# The influence of Temperature and Relative Humidity on the Development and Survival of Laboratory-Reared *Anopheles arabiensis* in Ethiopia

**DOI:** 10.1101/2025.04.30.651405

**Authors:** Temesgen Erena, Ademe kebede, Chernet Tuge Deressa, Delenasaw Yewhalaw, Eba Alemayehu Simma

## Abstract

Malaria, a major vector-borne disease transmitted by female Anopheles mosquitoes, remains a critical public health challenge in tropical and subtropical regions. Climate change influences vector distribution, abundance, and disease transmission patterns. This study examined the influence of temperature and humidity on the development, abundance, and survival of *Anopheles arabiensis*, a key malaria vector in Ethiopia. Laboratory reared susceptible *An. arabiensis* strain mosquitoes were reared under varying conditions, with 400 eggs exposed to temperatures of 14.5–34.35°C and relative humidity (RH) levels of 64–84% and compared with the ones that reared under standard insectary conditions. Development and survival rates were analyzed across 20 temperature-humidity regimes using SPSS 26, with linear regression, OLS, and quadratic regression models. Results showed optimal egg hatchability (80%) at 25.3–34.35°C and 64–68% RH. Adult emergence peaked (80%) at 28.37–29.65°C and 70% RH, but declined significantly below 16°C and 64% RH. The longest larval development (13 days) occurred at 14.55–15.62°C and 83–86% RH, while adult survival reached 25 days at 23.65–24.72°C and 68–69% RH. These findings highlight the profound influence of temperature and humidity on *An. arabiensis* life stages, emphasizing their role in mosquito population dynamics. Further research is needed to explore how these factors affect malaria parasite development within vectors, offering insights to improve malaria control strategies.

## Background

Malaria, a major vector-borne disease transmitted by female *Anopheles* mosquitoes, remains a critical public health challenge in tropical and subtropical regions. According to the World Health Organization (2023), an estimated 249 million malaria cases occurred in 2022 across 85 endemic countries, with sub-Saharan Africa bearing the highest burden, 93.6% of cases and 95.4% of deaths. Despite global efforts to combat the disease, malaria persists in sub-Saharan Africa, underscoring the region’s ongoing challenges in achieving effective control.

In Ethiopia, 47 *Anopheles* species have been documented, with *Anopheles arabiensis* identified as the primary malaria vector [1]. Secondary vectors include *An. funestus*, *An. pharoensis*, and *An. nili*. The recent invasion of *An. stephensi*, a major urban malaria vector, has further exacerbated the malaria burden in Africa [2]. Climate change significantly influences vector-borne diseases by altering mosquito distribution and expanding their geographical range [3-4]. Climate variability, characterized by short-term weather fluctuations, and climate change, defined as long-term shifts in weather patterns, both play critical roles in shaping mosquito ecology and malaria transmission dynamics. Variations in temperature, precipitation, and humidity directly influence mosquito survival, reproduction rates, and the development of the *Plasmodium* parasite within mosquito vectors. For instance, increased temperatures can accelerate the life cycles of both mosquitoes and parasites, potentially leading to higher transmission rates. Additionally, changes in rainfall patterns can create new breeding sites or eliminate existing ones, thereby altering mosquito population densities and distribution. These climatic factors collectively impact the geographical spread and seasonality of malaria, posing significant challenges to control and eradication efforts [5], [6]. Key indicators of malaria transmission include mosquito developmental traits, gonotrophic cycles, egg hatchability, and longevity, all influencing vectorial capacity. Temperature-dependent variations in gonotrophic cycle length affect transmission, as seen in *An. pseudopunctipennis* in Bolivia. Longevity is crucial, as only older mosquitoes transmit malaria after the extrinsic incubation period, with survival rates highly sensitive to climatic factors, especially temperature [7-8].

Mosquito growth and population dynamics are influenced by both biotic and abiotic factors, with temperature, humidity, and precipitation being the most critical abiotic drivers. These weather conditions significantly affect mosquito life cycles, population abundance, and disease transmission dynamics [9]. Moderate rainfall creates favorable breeding conditions for mosquitoes, while heavy rainfall can wash away breeding sites, potentially reducing malaria incidence. High rainfall has been associated with lower malaria transmission, possibly due to the flushing of mosquito habitats. Intense precipitation can disrupt mosquito populations by washing away breeding sites, thereby reducing malaria transmission in the short term [10-11].

Temperature is the most influential abiotic factor, affecting mosquito behaviour, growth, survival, dispersal, and reproduction. It impacts all juvenile stages: egg, larval, and pupal development [12-13]. Humidity also plays a role in egg hatchability [14], and rising temperatures can negatively affect larval survival, development time, and pupation success [15]. However, predicting population dynamics requires more than understanding these traits; adult production capacity and sex ratios are often overlooked in studies. While some research has explored temperature effects on sex ratios in *Aedes* mosquitoes [16-17], similar studies on *Anopheles* are scarce, particularly for Ethiopian *An. arabiensis* strains. This study addresses this gap by examining the combined effects of temperature and relative humidity on the development, survival, and sex ratios of *An. arabiensis*.

The broader implications of climate change on malaria incidence and mortality remain poorly understood. Rising temperatures could enable malaria to spread to previously unaffected areas, such as high-altitude regions above 2,000 meters [18]. Few studies have comprehensively explored the effects of temperature and humidity on mosquito life table parameters [19], highlighting the need for further research. This study investigates how temperature and relative humidity influence the development, survival, and abundance of *An. arabiensis* across all life stages. By analysing these factors, we aim to provide insights into the environmental drivers of mosquito population dynamics, contributing to more effective malaria control strategies.

## METHODS

### Descriptions of study area

The study was conducted from September 2023 to April 2024 at the Tropical and Infectious Diseases Research Centre (TIDRC) of Jimma University, located in Sekoru, Oromia Region (7.922305°N, 37.395320°E). Situated 246 km southwest of Addis Ababa at an altitude of 1,684 meters above sea level, the research center lies in a region characterized by distinct climatic patterns. The area experiences one dry season, lasting from November to March, and two rainy seasons: a long rainy season from June to September, peaking in July and August, and a short rainy season from April to May. The mean annual rainfall is 1,940.5 mm, with heavy rainfall occurring from June to August and lighter rains from September to December. Temperatures in Sekoru District range from 13°C to 20°C annually, with the highest temperatures typically recorded in August and the lowest in February (Ethiopian National Meteorological Agency, unpublished report).

### Mosquitoes rearing

The study involved rearing *An. arabiensis* mosquitoes under controlled environmental conditions, simulating a range of temperatures and relative humidity (RH) levels. A total of 400 eggs were obtained from the JU TIDRC insectary and reared to adulthood. The mosquitoes were exposed to temperatures ranging from 14.55°C to 34.35°C and RH levels between 64% and 86%. For the experimental setup, 8,000 eggs were distributed across 20 treatments, each with four replicates, while 3,200 eggs were allocated to the control group under eight treatments. After a 24-hour period, the eggs were carefully washed and transferred into hatching trays. Throughout the larval stage, the mosquitoes were provided with adequate nourishment, and their developmental progress was monitored and recorded until pupation. Once pupation occurred, the pupae were transferred into cages, and adult emergence was closely monitored until the end of their lifespan. Detailed records were maintained to document key developmental parameters, including egg hatching duration, as well as the survival and mortality rates of eggs, larvae, pupae, and adult mosquitoes under each temperature and humidity condition.

A control group was established under constant temperature and humidity conditions, following the same procedures as described earlier, to serve as a reference for the experiment. The insectary environment was carefully maintained at standardized conditions to ensure optimal growth and development across all mosquito life stages. Adult mosquitoes were reared at a stable temperature of 25±2°C and a relative humidity of 80±10%, while larvae and pupae were nurtured at 31±2°C with the same relative humidity level (80±10%). Throughout the study, records were kept, documenting critical parameters such as the duration of egg hatching, as well as the survival and mortality rates of eggs, larvae, pupae, and adult mosquitoes. This controlled environment provided a reliable baseline, allowing for precise comparisons with results obtained under varying simulated conditions.

### Collection of Larvae and pupae

Four replicates of larval instars (1st, 2nd, 3rd, and 4th) were reared in plastic containers measuring 35 × 25 × 10 cm. The larvae in each container were fed a pinch (approximately the amount held between the tip of the index finger and thumb) of finely ground Tetrafin fish food or cat food to ensure balanced and sufficient nutrition for optimal development. Daily observations were conducted, recording the duration of survivorship between each developmental stage (L1–L2, L2–L3, L3–L4, and L4 to pupae), the number of larvae, and the corresponding temperature and humidity conditions. Larval mortality rates for each instar were also recorded.

Pupae were collected twice daily, in the morning and afternoon, using a plastic dropper. The collected pupae were placed in disposable 150 ml plastic cups, with an average of 100 pupae per cup filled to two-thirds with water. These cups were then transferred into metal-framed cages (30 × 30 cm) covered with netting, where the pupae developed into adult mosquitoes. The time taken for larvae to develop into pupae, the number of pupae, and the duration from pupal emergence to adulthood were recorded. Additionally, pupal and adult mortality rates were recorded throughout the process.

### Adults Emergence and development

After pupation, adult mosquitoes emerged and were housed in clean metal cages placed in the adult room. The cages were labelled to indicate the replication type and collection date. The mosquitoes were provided with a sugar solution (one part sugar to nine parts dechlorinated water) for nourishment. Cotton balls were soaked in the sugar solution until fully saturated and then placed on inverted plastic cups inside each cage. Female adult mosquitoes were blood-fed to support egg development. For blood feeding, the belly of a rabbit was shaved and secured to a wooden frame, which was then placed on top of the cage. The females fed through the cage netting until their abdomens became visibly red and swollen with blood. Daily mortality and survivorship rates were recorded to monitor the adult mosquito population.

### Data analysis

Entomological and climatological data, including the number of immature and adult mosquitoes developed, the duration of each life cycle stage, and daily mean temperature and relative humidity, were recorded and calculated over a seven-month period (September 2023 to April 2024). The data were entered into Microsoft Excel and subsequently exported to the Statistical Package for the Social Sciences (SPSS) version 26 for analysis. A significance level of p < 0.05 was set for all statistical tests. The number of immature and adult mosquitoes developed was analyzed using linear regression, while Ordinary Least Squares (OLS) regression was employed to examine the relationship between temperature and relative humidity factors. The mean differences in the duration of each life cycle stage were assessed using a two-way ANOVA. Additionally, future malaria vector population trends were predicted using quadratic and polynomial linear regression models.

## RESULTS AND DISCUSSION

### Results

From September 2023 to March 2024, we observed that a fluctuation in climate (temperatures and relative humidity) significantly impacted on the development and survivorship of *An. arabiensis* strain. Temperature fluctuations and changes in relative humidity significantly influenced the production of first instar larvae. The highest production of L1 larvae (93%) occurred at higher temperatures (33.35℃) and lower relative humidity (68%). Optimal conditions for egg hatchability were observed at temperatures ranging from 26 to 34 ℃ and relative humidity levels between 68 and 70%. In contrast, at lower temperatures of 14.55℃ and 15.62℃ combined with higher relative humidity levels of 86% and 83%, respectively, larval production was significantly reduced, with only 33% of eggs successfully hatching (Fig. 1). The mean number of hatched eggs (L1) as a function of temperature and relative humidity showed statistically significant effects (*P* < 0.001).

**Fig. 1:**
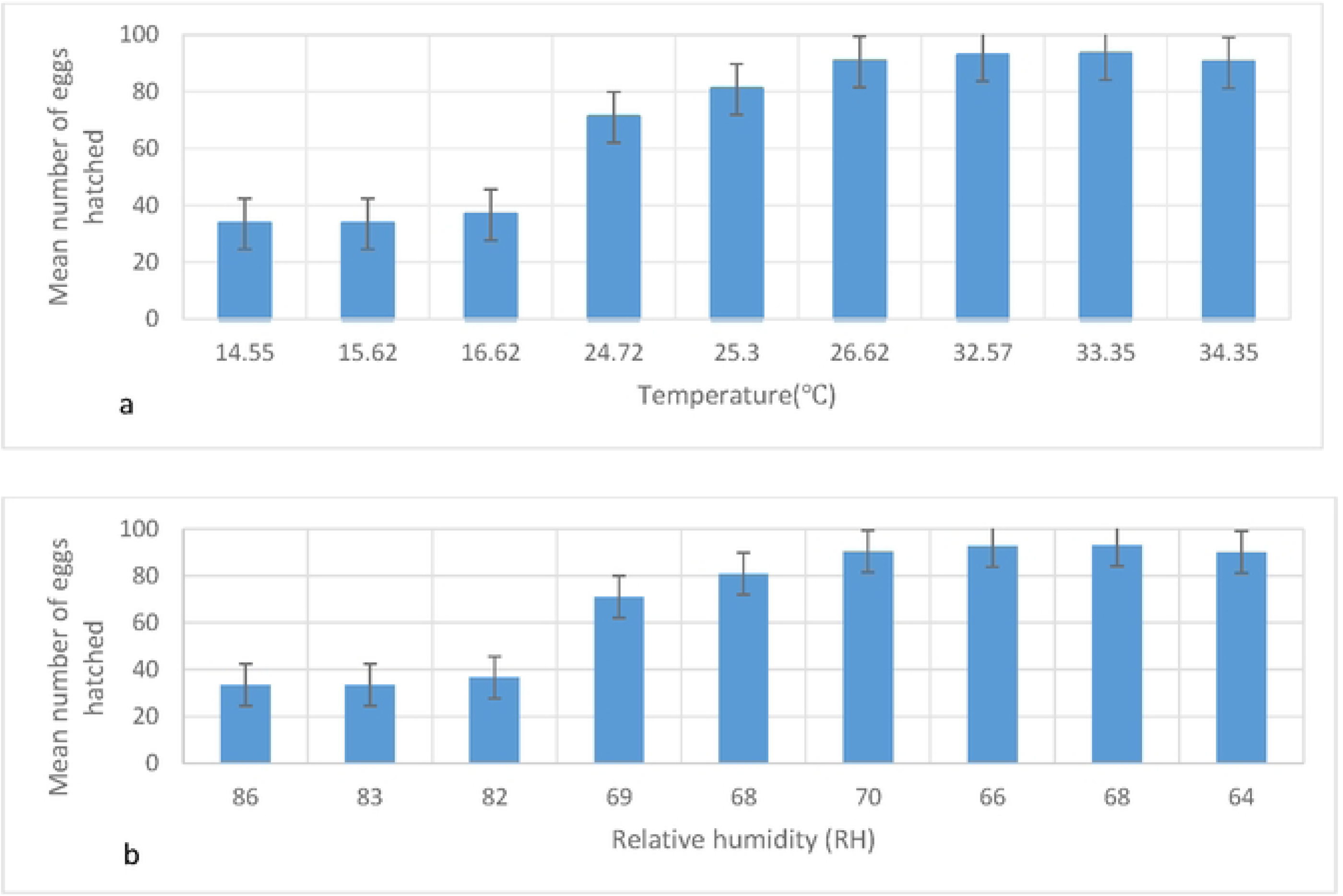
Mean eggs hatchability rate of *An. arabiensis* at different temperature (a) and relative humidity levels (b)

An increase in temperature and a decrease in relative humidity were associated with a higher frequency of second ecdysis. The optimal conditions for the development of second instar larvae (L2) were observed at temperatures ranging from 26-34 ℃ and relative humidity (RH) between 64 and 70%.The highest number of L2 larvae occurred at 32.57℃ and 66% RH, while the lowest numbers were recorded at 15.62℃, 83% RH (Fig. 2). Both temperature and relative humidity were found to have a statistically significant impact on L2 development (*P* < 0.001), with specific significance level (P=0.002).

**Fig. 2:**
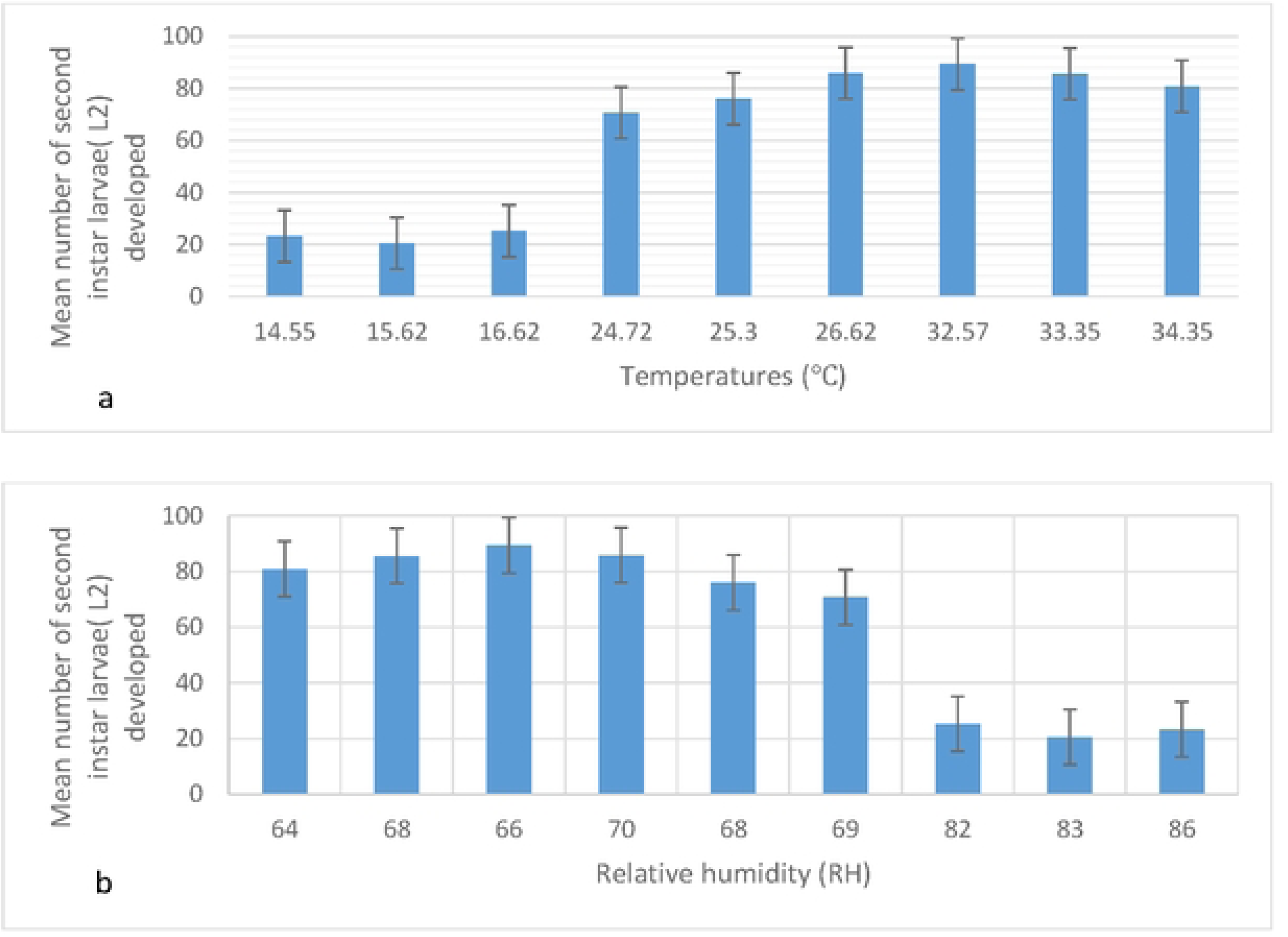
Mean number of second instar larvae (L2) developed at different temperature (a) and relative humidity levels (b)

The mean development rate from second instar (L2) to third instar (L3) increased with rising temperature and relative humidity. The optimum temperature for L3 development were observed at temperatures ranging from 26 to 34 ℃ and relative humidity (RH) levels between 64 and 70%. The highest number of third instar larvae (L3) was produced at 32.57℃ and 66% RH, while the lower number of L3 larvae was recorded at 15.62 ℃ and 83% RH (Fig. 3).

**Fig. 3:**
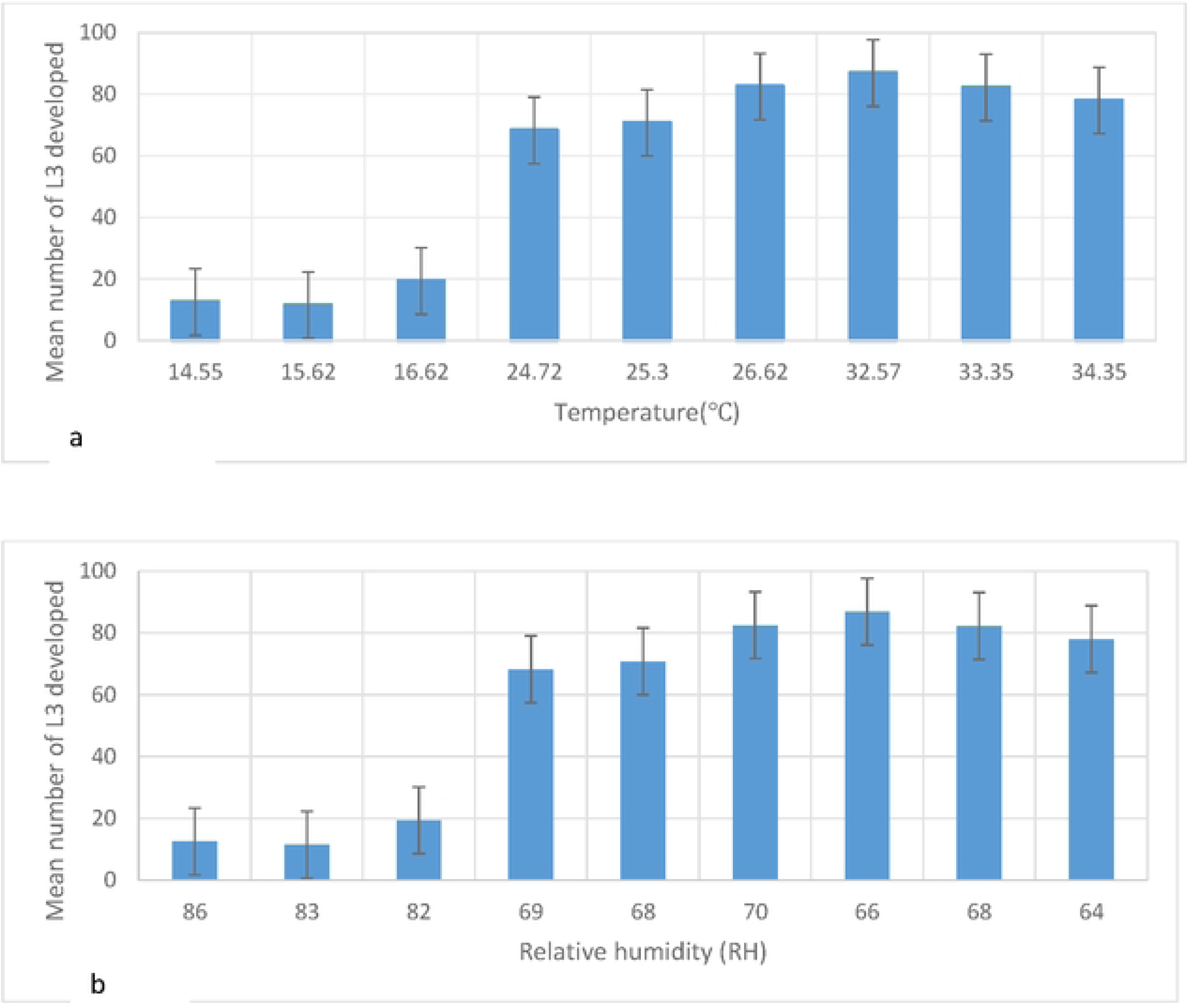
Mean number of 3^rd^ instar larvae developed at different temperature (a) and relative humidity levels (b)

The mean development rate of fourth instar (L4) increased with rising temperatures and decreasing relative humidity (Fig. 4). The highest L4 count was recorded at 28.37℃ and 70% RH, while the lowest occurred at 15.62℃ and 83% RH. The optimal temperature and relative humidity ranges for L4 development were 26-34 ℃ and 70-64% respectively. Both temperature and relative humidity had a significant effect (*P* < 0.001, with P=0.002).

**Fig. 4:**
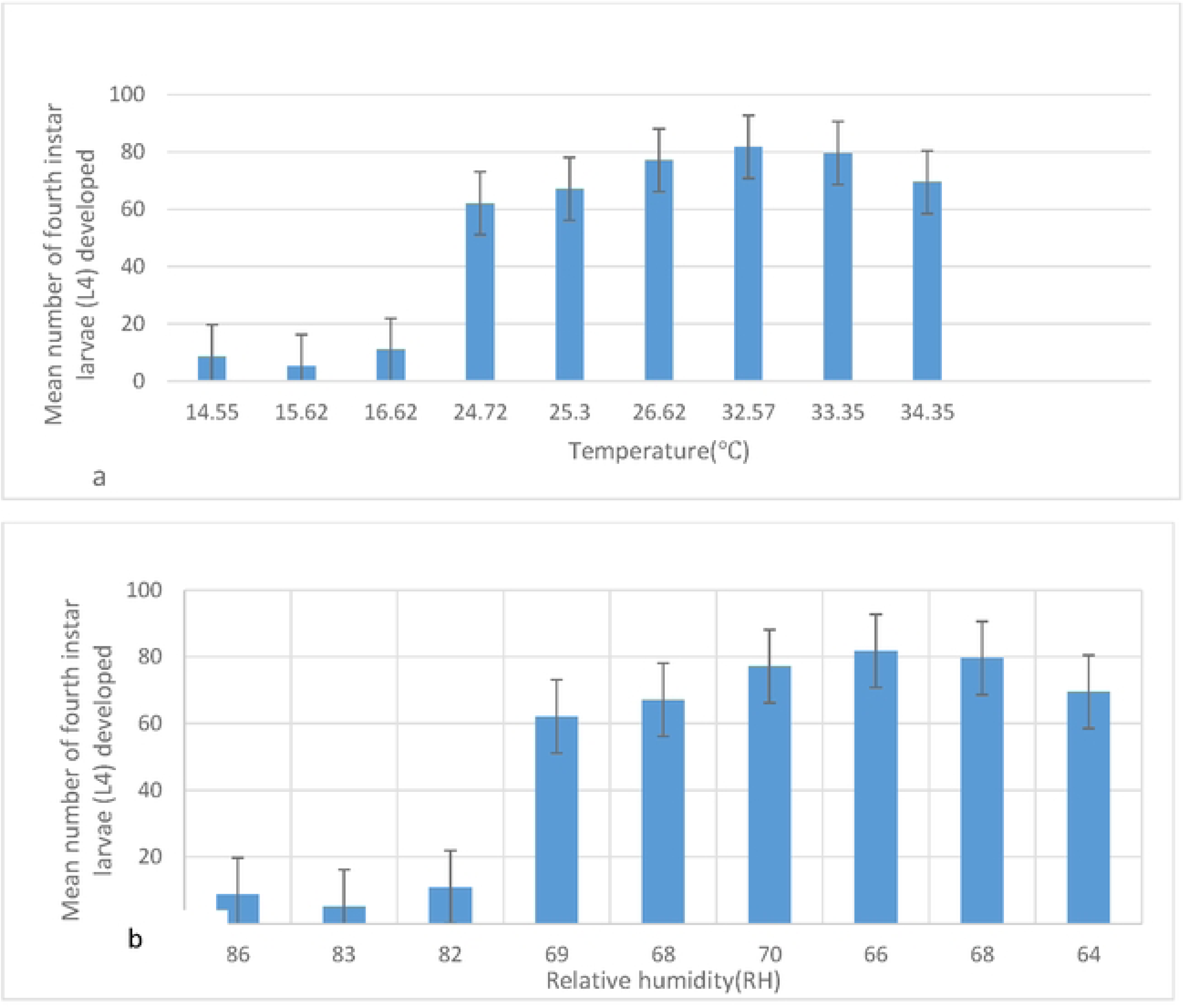
Mean number of fourth instar larvae (L4) developed at different temperature (a) and relative humidity levels (b)

Increasing temperatures were found to accelerate pupal development, while decreasing the relative humidity slowed it down (Fig. 5). The optimal conditions for pupation were observed at temperatures ranging from 32 to 34℃ and relative humidity levels between 64-68%.

**Fig. 5:**
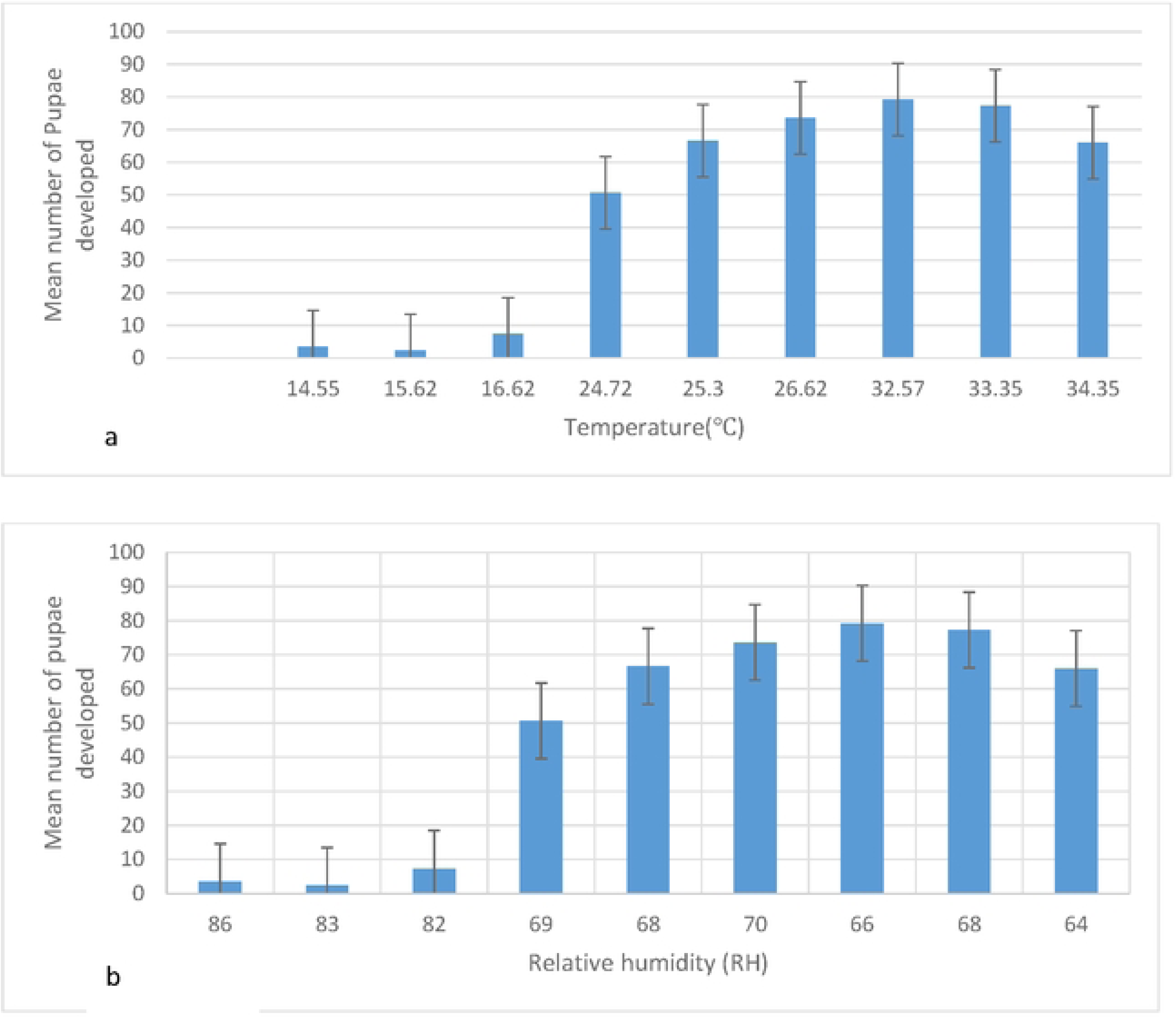
Pupation at different temperature (a) and relative humidity levels (b)

As temperature increases and relative humidity decreases, the number of developing adults also increases. However, beyond 31℃, below 68% RH, adult development rates decline. The optimal conditions for adult development were observed at temperatures ranging from 24 to26℃ and RH levels between 68 and 70%. The peak adult development rate (60%) observed at temperatures of 25℃ - 26℃ and RH levels 68-70% (Fig. 6).

**Fig. 6:**
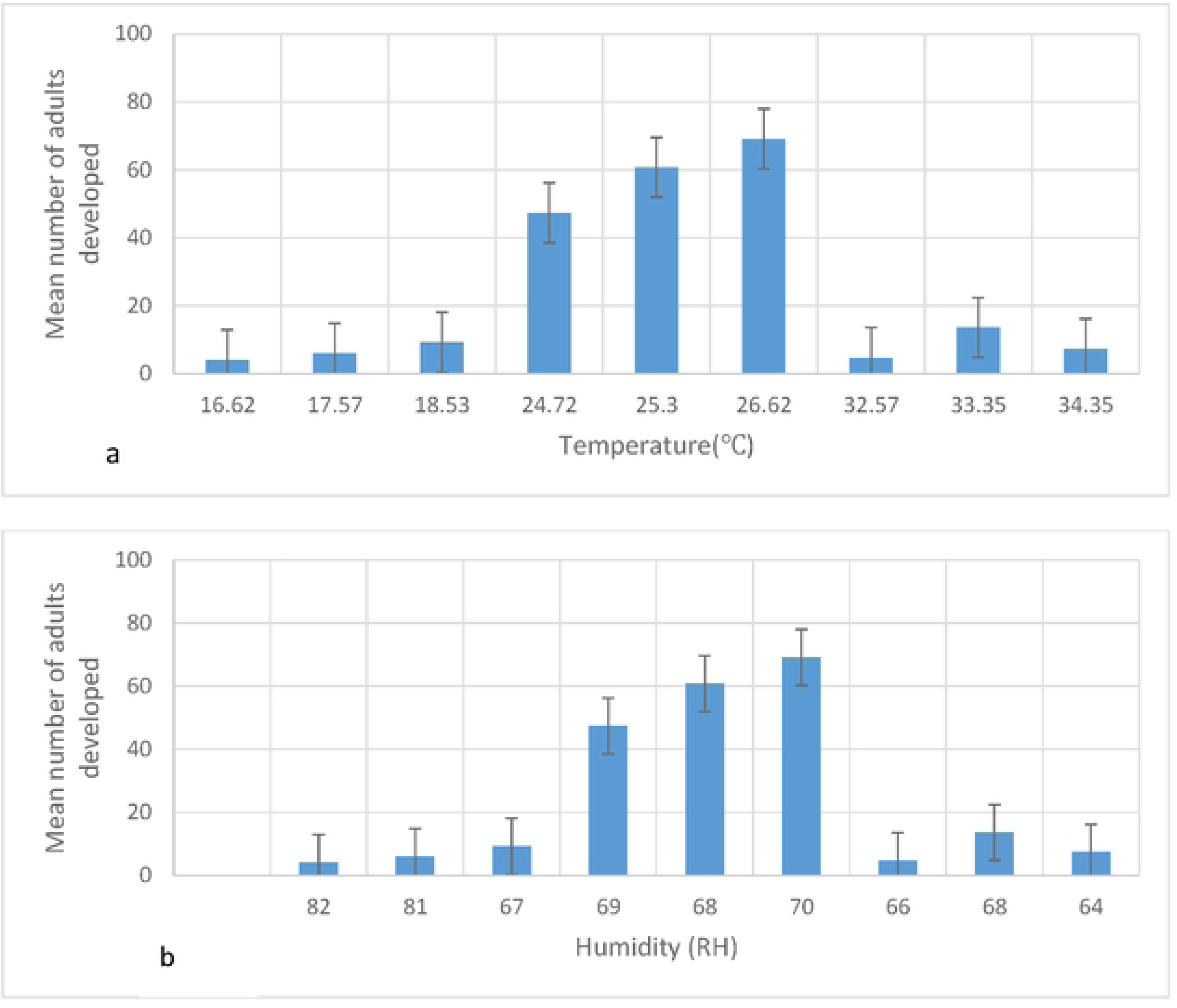
Adults development at different temperature (a) and relative humidity levels (b)

The number of days required for egg hatching increased with higher relative humidity and lower temperatures. The optimal conditions for egg hatching were observed at temperatures between 14 to16℃ and relative humidity 82-86%. The maximum time required for egg hatching was 11 days at 14.55℃ and 86% RH, while the minimum time of two days was recorded at 34.35℃ and 64% RH (Fig. 7). However, the relationship between these conditions and hatching time was not statistically significant (P=0.207).

**Fig. 7:**
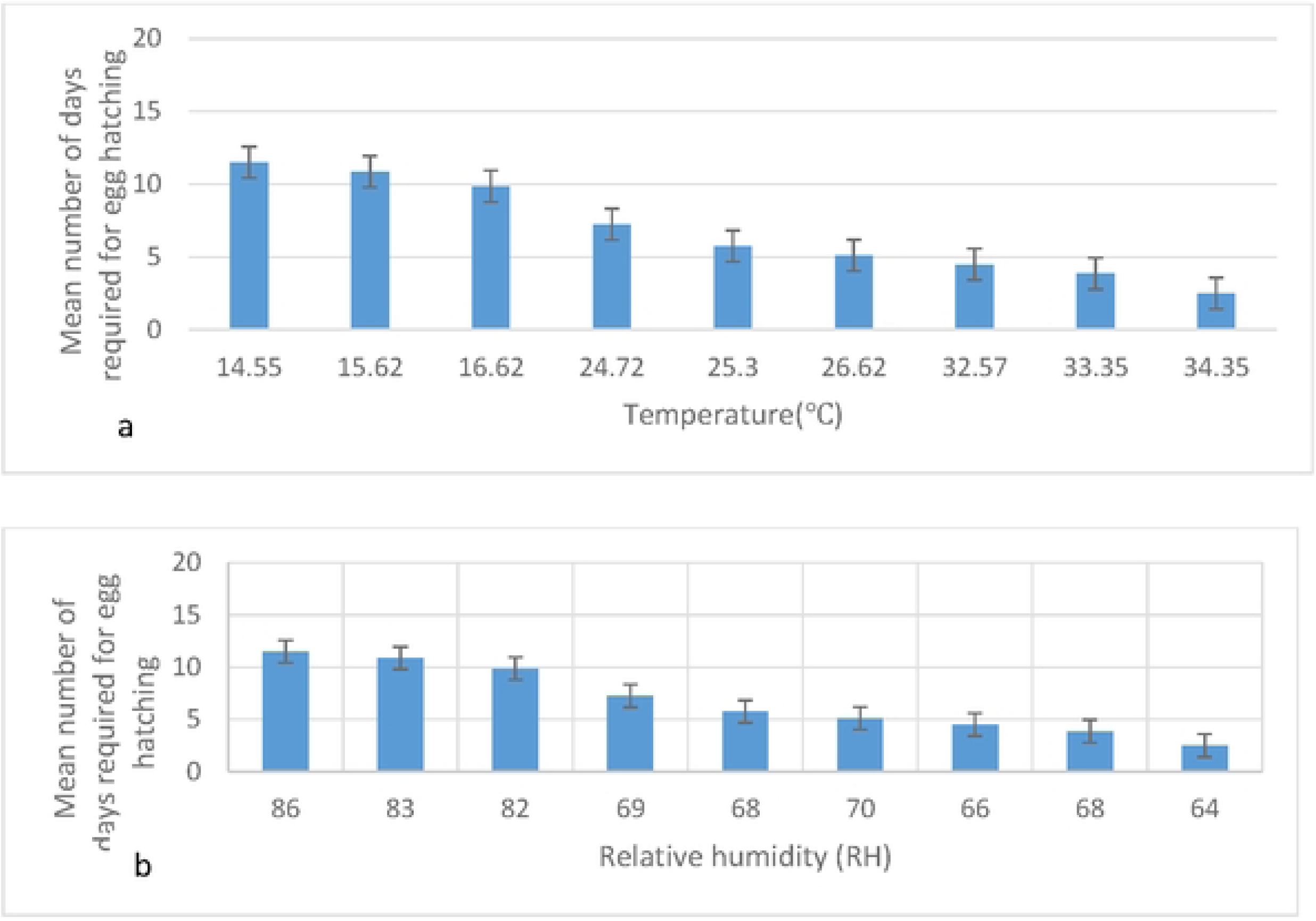
Number of days required for egg hatching at different temperatures (a) and relative humidity levels (b)

Increasing relative humidity extended the number of days required for the development of second instar larvae (L2), while decreasing temperatures also prolonged the L2 developmental period. The maximum days required for second instar larvae (L2) development was 13 days, whereas the minimum time of 4 days was observed at 34.35℃ and 64% RH (Fig. 8). On average, the development from first instar larvae (L1) to second instar larvae (L2) took between 4 and13 days. The optimal conditions for L2 development were found to be temperatures ranging from14-25℃ and relative humidity levels between 71-86%.

**Fig. 8:**
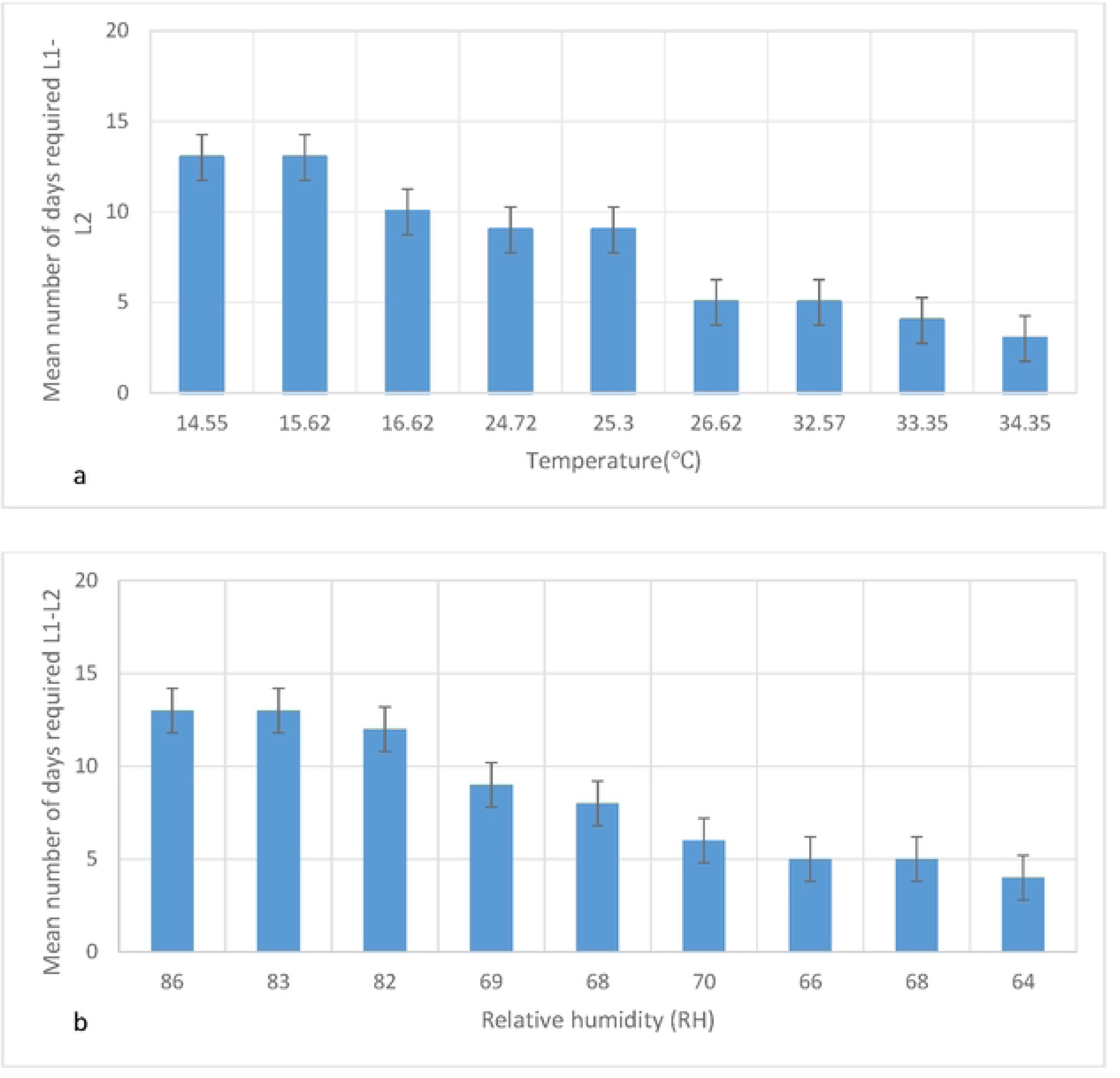
Number of days required to develop from L1 to L2 at different temperature (a) and relative humidity levels (b)

Higher relative humidity prolonged the number of days required for the development of third instar larvae (L3), while lower temperature also increased the duration of third instar larvae (L3) development (Fig. 9). The optimal conditions for L3 development were observed at temperatures ranging from 14 to26℃ and relative humidity levels between 68-86%. However, the effects of both temperature and relative humidity on L3 development were statistically insignificant (P=0.119). On average, the development from L2 to L3 took between 2 and11days.

**Fig. 9:**
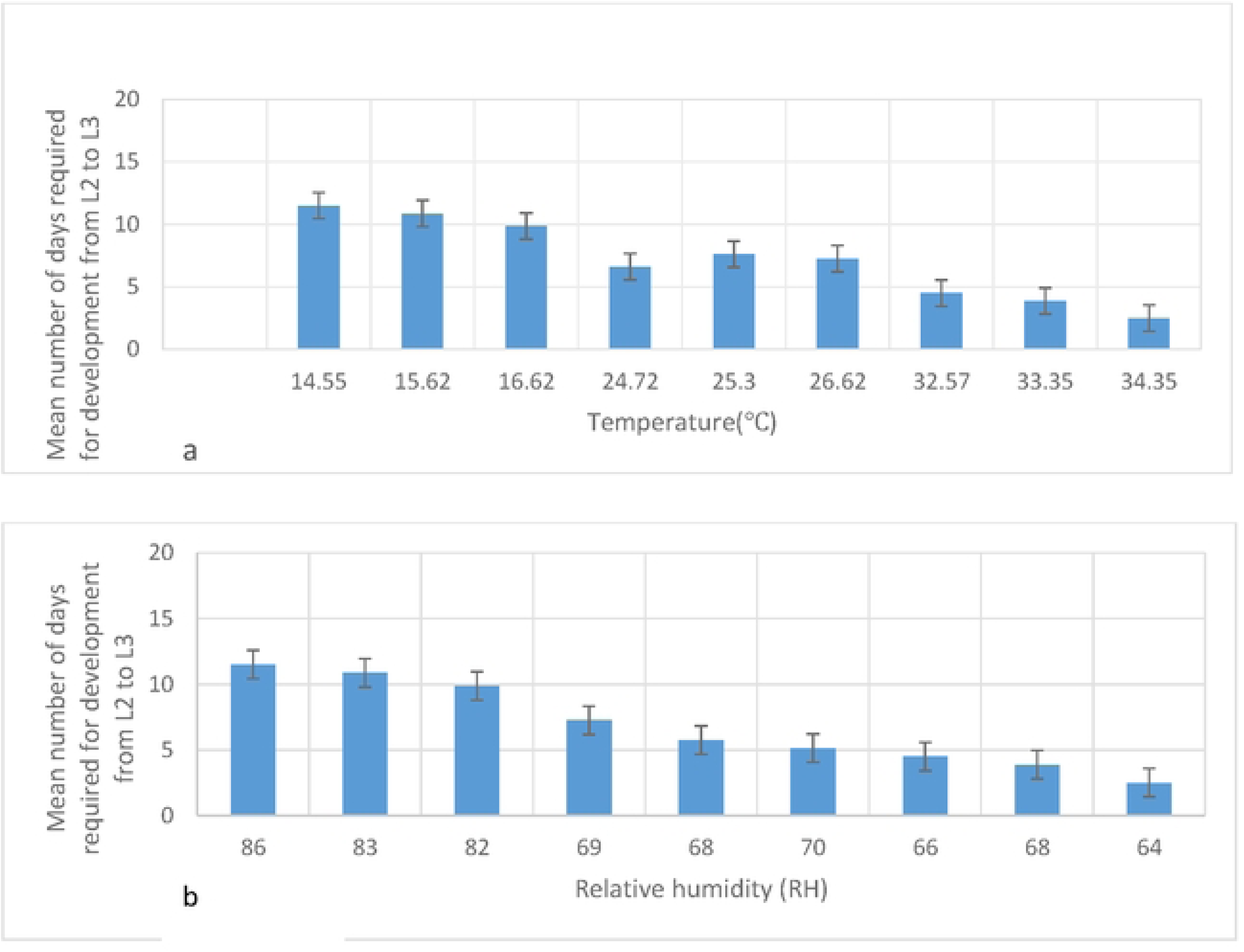
Number of days required to develop from L2 to L3 at different temperature (a) and relative humidity (b)

Higher relative humidity extended the number of days required for the development to fourth instar larvae (L4), while lower temperatures also increased the duration of L4 development. The longest development period for L4 lasting six day, was observed at 15.62℃ and 83% RH. In contrast, the shortest development period, requiring only one day, was recorded at 34.35°C and 64% relative humidity (Fig. 10). On average, the development from third instar larvae (L3) to L4 took between 2 and 6 days.

**Fig. 10:**
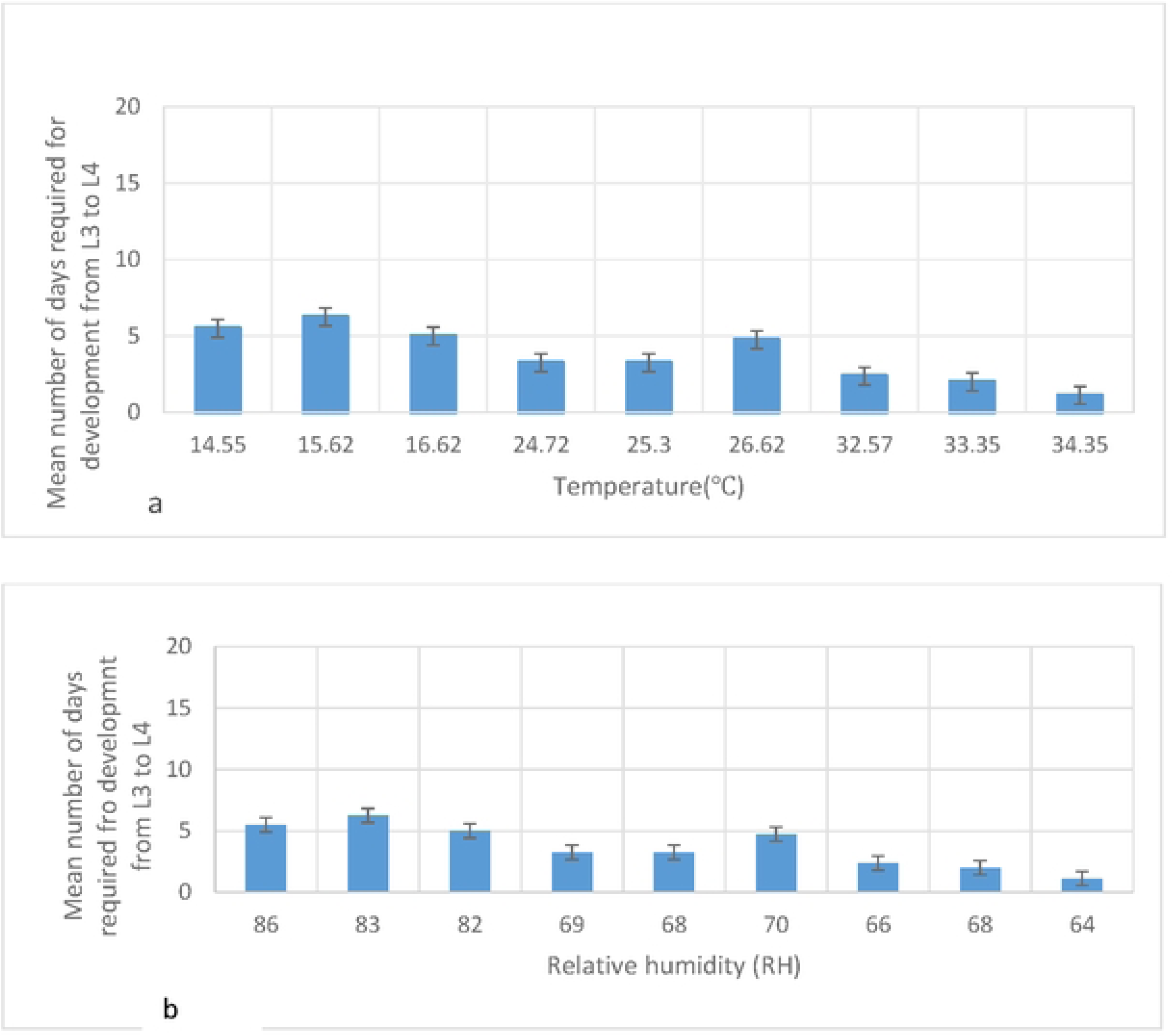
Number of days required for larval stage 4 developments at different temperature (a) and relative humidity (b)

Pupation rates were significantly influenced by environmental conditions, with higher rates observed at increased relative humidity and lower rates at elevated temperatures. The average duration for L4 development to pupae ranged from 3 to 7 days. The maximum number of days required for pupation was seven, recorded at 15.62℃ and 83% RH (Fig. 11). Both temperature and relative humidity were found to have statistically significant impact on pupation (*P* < 0.001, P=0.000).

**Fig. 11:**
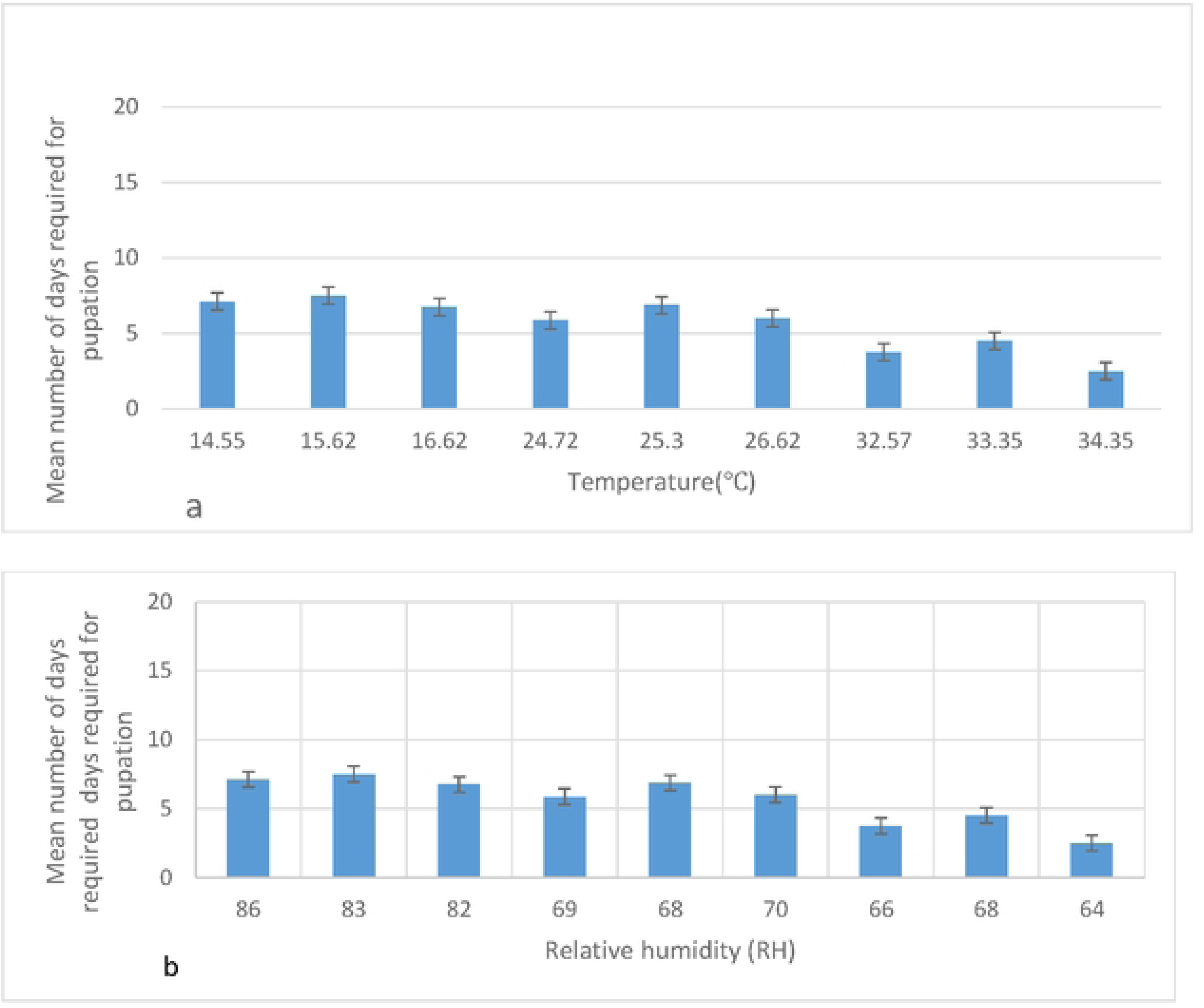
Number of days required for pupation at different temperature (a) and relative humidity (b)

The optimal conditions for adult were observed at temperatures ranging from 16-20℃, and RH levels82-74% (Fig. 12). Additionally, the most favourable temperature and relative humidity for adult development were found to be 1725℃, 81-68% respectively.

**Fig. 12:**
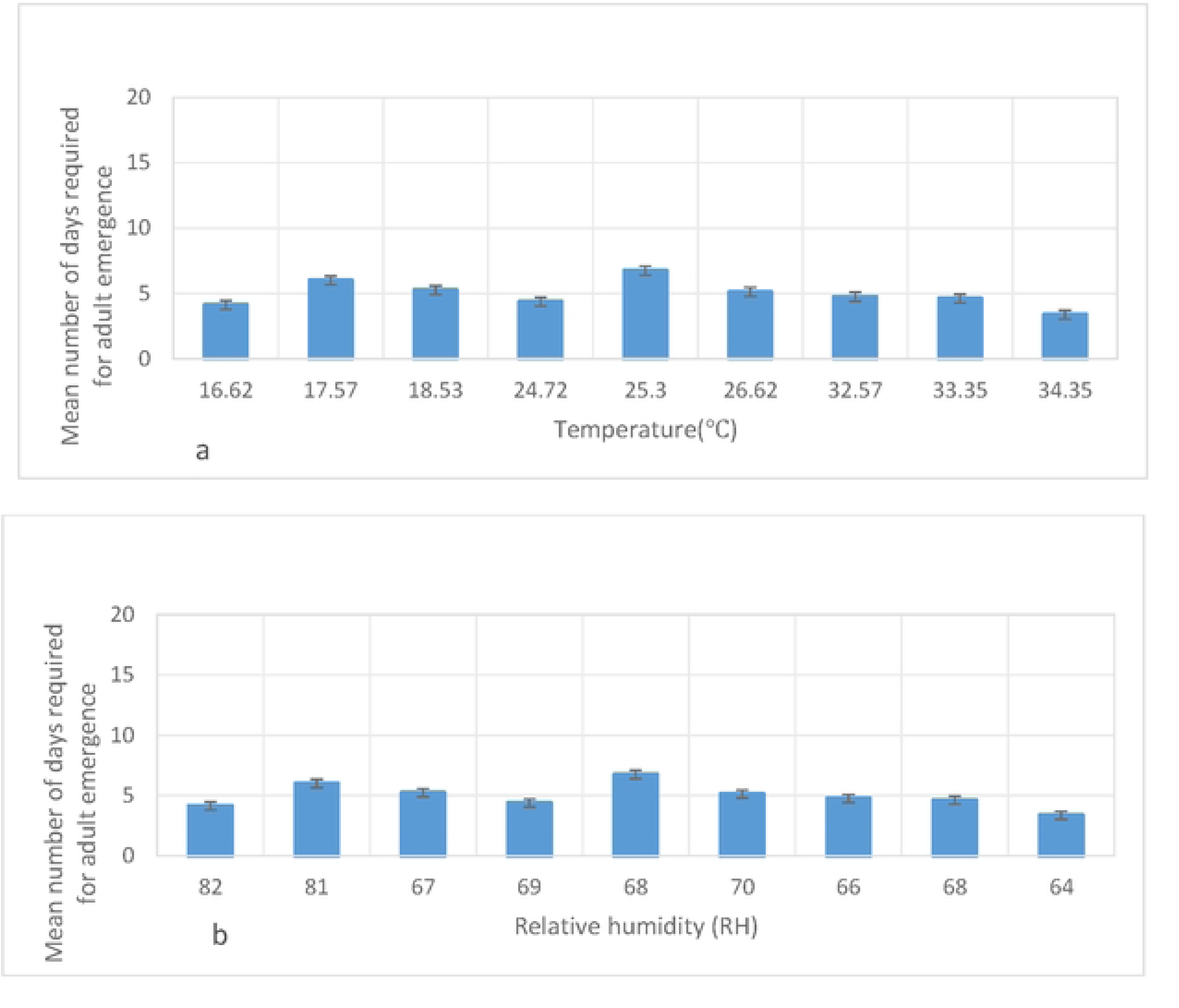
Number of days required for adult emergence at different temperature (a) and relative humidity (b)

At higher relative humidity and lower temperature (24℃ with 69% RH), the maximum adult survival time was observed to be 25 days. In contrast, the minimum survival time was 12 days at 34.35℃ and 64% RH (Fig. 13). The optimal conditions for adult survival were found to be within the temperature range of 24-26℃ and relative humidity of 69-70%. Both temperature and relative humidity were statistically significant factors influencing survival, with *P* < 0.001 (P=0.000).

**Fig. 13:**
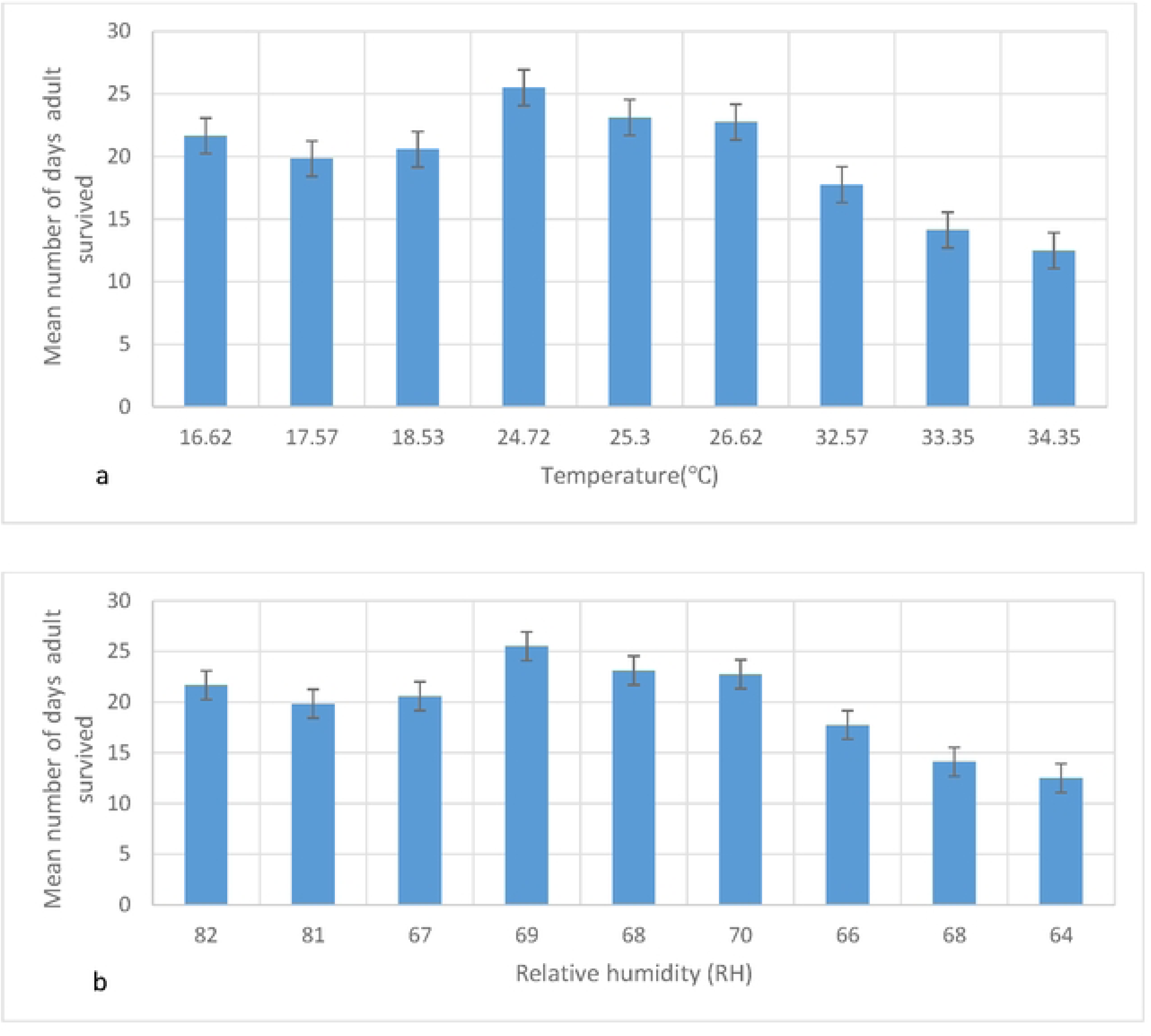
Mean number of days adult survived at different temperatures (a) and relative humidity levels (b)

At temperatures ranging from 16.62-24℃ with relative humidity between 82-69%, as well as higher temperatures of 32.5-34.35℃ with relative humidity of 64-66%, the production of male adult mosquitoes was significantly reduced(<10%). The lowest numbers of both male and female mosquitoes were observed at temperature below 20.6 ℃ with 74% RH. A substantial proportion of larvae and pupae failed to reach adulthood under these conditions, leading to decline adult mosquito populations. This was particularly evident under extreme environmental conditions, including high temperatures with low relative humidity, as well as low temperatures with high relative humidity (Fig. 14).

**Fig. 14:**
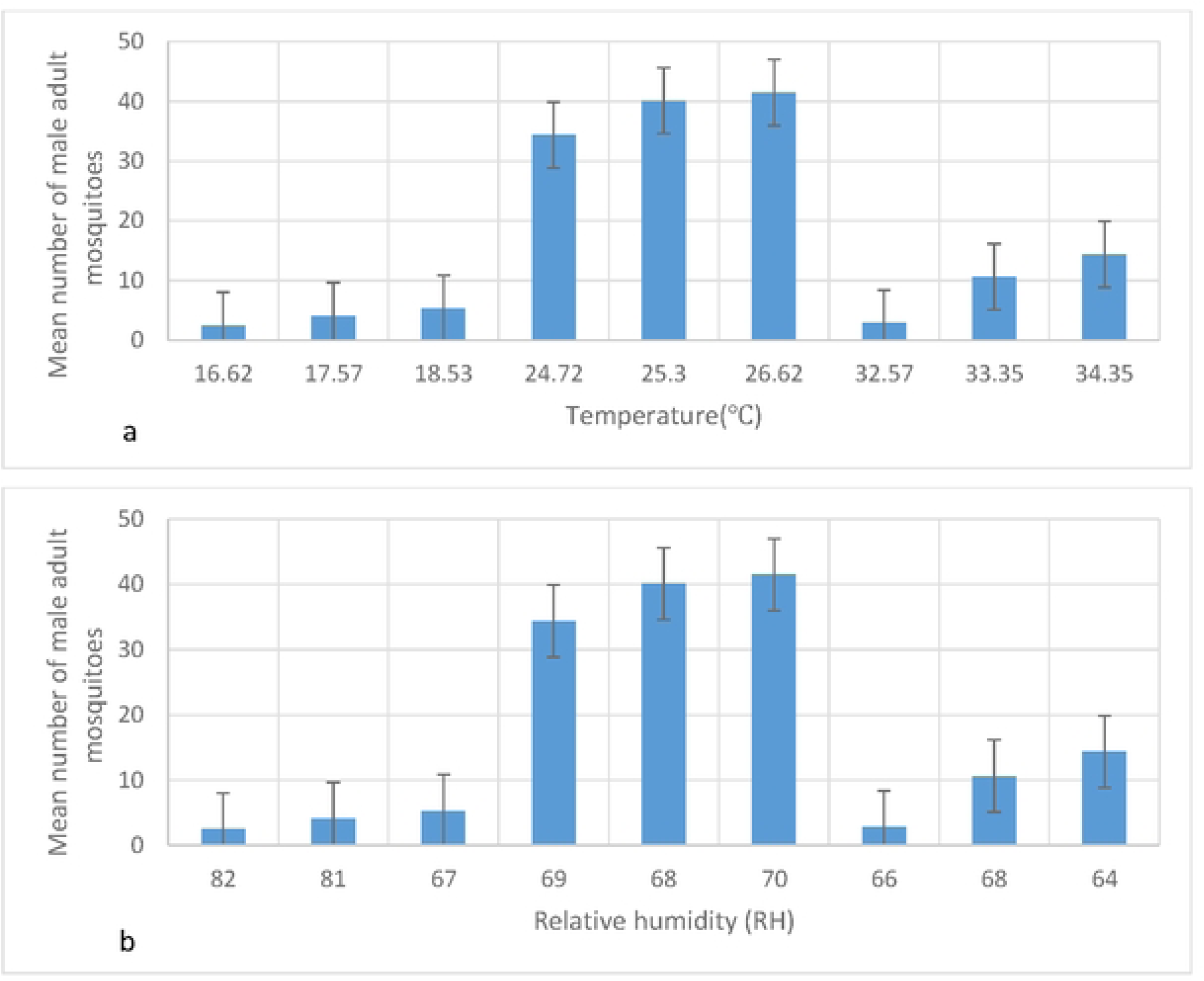
Mean number of male mosquitoes at different temperatures (a) and relative humidity levels (b)

The production of female adult mosquitoes was significantly reduced under certain environmental conditions (at lower temperatures and higher humidity (16.62-24℃, 82-69% RH), fewer than 20% of females reached adulthood survived, while even fewer (<10%) survived at higher temperatures of (32.5-34.35℃ and higher humidity of 66-64% RH (Fig. 15). The optimal conditions for female mosquito production were observed at temperatures of 22-31℃ and relative humidity of 68-70%. However, these factors were not statistically significant, as indicated by a P-value of 0.111.

**Fig. 15:**
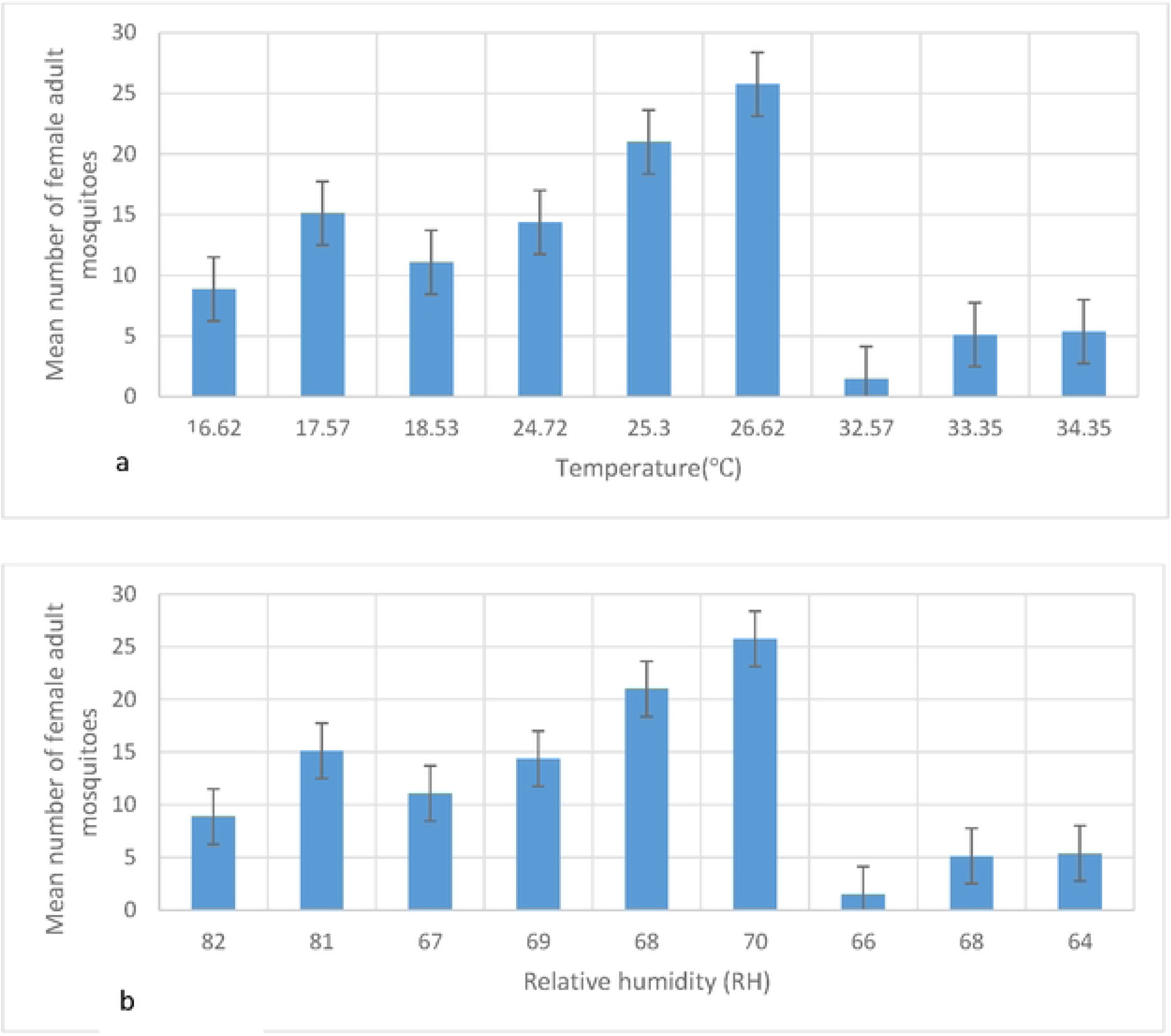
Mean number of female mosquitoes at different temperatures (a) and relative humidity levels (b)

## Discussion

The study highlights the significant influence of temperature and relative humidity on the life cycle dynamics of *An. arabiensis* mosquitoes. The findings reveal a linear relationship between these environmental factors and the development and survival of both immature stages (larvae and pupae) and adult populations, ultimately affecting their developmental time and overall survival [13,16,19]. These insights are critical for understanding how variations in climatic conditions can shape malaria transmission dynamics, providing valuable information for predicting and mitigating the impact of climate change on vector-borne diseases..

The lower development threshold (LDT) for *An. arabiensis* larvae was identified as approximately 14.55°C, while the upper development threshold (UDT) was around 34.35°C. Egg hatching rates increased with rising temperatures as temperature increased and decreasing relative humidity, leading to higher numbers of 1st instar larvae under these conditions. However, higher temperatures significantly reduced larval survival time, with no larvae surviving above 35°C as corroborated by previous studies [8,13, 20,21].

Pupation was significantly influenced by environmental factors, with higher relative humidity promoting the process and elevated temperatures hindering it. The longest pupation duration (7 days) was observed at 15.62℃ and 83% relative humidity, while the shortest duration (2 days) occurred at 34.45℃ and 64% relative humidity. This suggests that optimal pupation conditions require a delicate balance between temperature and humidity levels. These findings contrasts with those of [22], who reported that higher temperatures did not significantly affect the survival of immature stages of *An. arabiensis* and *An. funestus.* This discrepancy indicates that the impact of temperature on development may vary across species or under different environmental conditions, emphasizing the need for further research to better understand these dynamics.

Our study demonstrated that Anopheles mosquito survival and development are highly temperature-dependent. Optimal adult emergence (80%) occurred within 28.37–29.65°C at 70% relative humidity, while temperatures below 16°C, especially with humidity above 82%, entirely prevented development. A broader range (25–31°C) supported over 60% emergence, but outside this range, survival declined sharply, with minimal emergence below 20°C or above 31°C. Additionally, temperatures above 29°C increased adult mortality, reducing reproductive output.

These findings align with studies showing that *Anopheles* development peaks within a narrow temperature range, with extreme temperatures hindering survival [22-23]. Similar patterns have been observed in *An. gambiae*, where optimal emergence occurs around 27–30°C, with reduced survival beyond this range [13]. However, some studies suggest *An. funestus* may tolerate slightly broader temperature fluctuations [24]. These variations underscore species-specific thermal sensitivity and the importance of local environmental conditions in shaping mosquito population dynamics.

The longevity of adult *Anopheles* mosquitoes is strongly influenced by environmental conditions, with high humidity and low temperatures negatively affecting survival. Our findings align with previous studies emphasizing the adverse effects of such conditions. Specifically, optimal survival was observed at higher temperatures (34.35°C) and lower humidity (64%), where the minimum survival time recorded was 12 days [19].

However, these results contrast with those of Barreaux et al. (2018), who reported that higher temperatures did not significantly affect the survival of *An. gambiae*. This discrepancy suggests that the impact of temperature on mosquito longevity may vary across species or under different environmental conditions, highlighting the need for further research to understand species-specific thermal tolerance and survival mechanisms [25].

Research conducted across various geographical regions, including Turkey [26], and South Africa [27], consistently demonstrates that rising temperatures reduce *Anopheles* mosquito survival rates. Elevated temperatures can disrupt mosquito ecology potentially intensifying the transmission of vector-borne diseases such as malaria [28-29]. Our findings emphasize the pivotal role of temperature and humidity in influencing mosquito development, survival, and reproductive success, underscoring their profound implications for malaria transmission dynamics in the face of climate change.

## CONCLUSION

Our findings highlight the critical role of temperature and humidity in shaping the life cycle and survival of *An. arabiensis* mosquitoes. These insights are essential for predicting malaria vector distribution and designing effective interventions in Ethiopia and similar environments. Extreme temperature and humidity levels can significantly impact mosquito lifespan—higher temperatures with lower humidity accelerate egg development and hatching, while lower temperatures with higher humidity extend survival but slow immature stage development and adult emergence.

Overall, warmer temperatures and lower humidity speed up mosquito development but shorten adult lifespan, whereas cooler temperatures and higher humidity prolong development. With climate change, *An. arabiensis* may expand into previously malaria-free regions, particularly within the optimal range of 16°C–34°C and 64%–82% humidity. This underscores the need for proactive malaria control measures as global climate patterns shift.

While this study extensively examines temperature and humidity effects on malaria vectors, future research should explore their influence on malaria parasite development within mosquitoes, including temperature and humidity thresholds. A deeper understanding of these interactions would enhance malaria control strategies by addressing both vector and parasite dynamics.

## Acknowledgements

The authors are grateful for the technical and logistical support provided by the Tropical and Infectious Diseases Research Center at Jimma University. They also express their thanks to the Department of Biology and College of Natural Sciences, Jimma University for the overall support during the study period.

## Corresponding authors

Correspondence to Eba Alemayehu Simma

## Author’s Contributions

CTD, EAS and DY conceived and designed the experiments; TE performed the experiments; TE, CTD, EAS and AK analyzed and interpreted the data; TE, EAS, CTD and DY wrote the manuscript. All authors read and approved the final manuscript.

## Consent for publication

Not applicable.

## Data availability

Data generated during the study are available from the corresponding authors up on request

